# A machine-learning heuristic to improve gene score prediction of polygenic traits

**DOI:** 10.1101/107409

**Authors:** Guillaume Paré, Shihong Mao, Wei Q. Deng

## Abstract

Machine-learning techniques have helped solve a broad range of prediction problems, yet are not widely used to build polygenic risk scores for the prediction of complex traits. We propose a novel heuristic based on machine-learning techniques (GraBLD) to boost the predictive performance of polygenic risk scores. Gradient boosted regression trees were first used to optimize the weights of SNPs included in the score, followed by a novel regional adjustment for linkage disequilibrium. A calibration set with sample size of ~200 individuals was sufficient for optimal performance. GraBLD yielded prediction *R^2^* of 0.239 and 0.082 using GIANT summary association statistics for height and BMI in the UK Biobank study (N=130K; 1.98M SNPs), explaining 46.9% and 32.7% of the overall polygenic variance, respectively. For diabetes status, the area under the receiver operating characteristic curve was 0.602 in the UK Biobank study using summary-level association statistics from the DIAGRAM consortium. GraBLD outperformed other polygenic score heuristics for the prediction of height (*p*<2.2x10^−16^) and BMI (*p*<1.57x10^−4^), and was equivalent to LDpred for diabetes. Results were independently validated in the Health and Retirement Study (*N*=8,292; 688,398 SNPs). Our report demonstrates the use of machine-learning techniques, coupled with summary-level data from large genome-wide meta-analyses to improve the prediction of polygenic traits.

## Introduction

The advent of precision medicine depends in large part on the availability of accurate and highly predictive polygenic risk scores. While progress has been made identifying genetic determinants of polygenic traits, the amount of phenotypic variance explained by polygenic risk scores derived from genome-wide significant associations remains modest. On the other hand, moderate to high narrow-sense heritability has been established for many human traits. It has been proposed that weak, yet undetected, associations underlie polygenic trait heritability^1^. Consistent with this hypothesis, polygenic risk scores that include both strongly and weakly associated variants are vastly superior than those including only genome-wide significant variants. For example, a recent study by Abraham *et al*. showed that a polygenic risk score incorporating 49,310 variants had a discrimination ability that was similar and complementary to the widely used clinical Framingham risk score for the prediction of coronary artery disease (CAD)^2^. Thus, there is a need for polygenic risk score methods that can integrate a large number of genetic variants.

The most popular heuristic for polygenic risk score is based on linkage disequilibrium (LD) pruning of SNPs, prioritizing the most significant associations up to an empirically determined *p*-value threshold, and pruning the remaining SNPs based on LD^3^. This “pruning and thresholding” (P+T) approach has the advantage of being simple and computationally efficient, but discards some information because of LD pruning. To remediate this issue, a novel method, LDpred, which uses LD information from an external reference panel, was recently proposed to infer the mean causal effect size using a Bayesian approach^4^. While the latter method has been shown to improve prediction *R^2^*, it depends on estimates of polygenic heritability and causal fraction, and can be sensitive to the misspecification of LD. We hypothesized that a further gain in prediction *R^2^* could be made by tuning the weights of SNPs included in polygenic risk scores using principles of machine-learning.

Machine-learning encompasses a wide-ranging class of algorithms widely used to solve complex prediction problems. It has proven particularly useful when prediction is dependent on the integration of a large number of predictors, including higher-order interactions, and when sizeable training datasets are available for model fitting. In particular, gradient boosted regression trees are powerful and versatile methods for continuous outcome prediction^5^, and thus, are ideal for updating the SNP weights in polygenic risk scores. Tree-based models partition the predictor space according to simple rules by identifying regions having the most homogeneous responses to predictors and fitting the mean response for observations in that region. Gradient boosting^6^ is an efficient algorithm that sequentially combines a large number of weakly predictive models to optimize performance.

We propose to leverage the large number of SNPs and the available summary-level statistics from genome-wide association studies (GWAS) to calibrate the weights of SNPs contributing to the polygenic risk score, adjusting for LD instead of pruning. Our heuristic, gradient boosted and LD adjusted (GraBLD; https://github.com/GMELab/GraBLD), involves two steps and uses the univariate regression coefficients from external meta-analysis^7-9^ summary association statistics as the starting point (see Figure 1 and Methods). First, the external univariate regression coefficients were updated with respect to a target population by the gradient boosted regression tree models. Second, the updated weights were corrected for LD to produce the final polygenic risk score.

**Figure 1:**
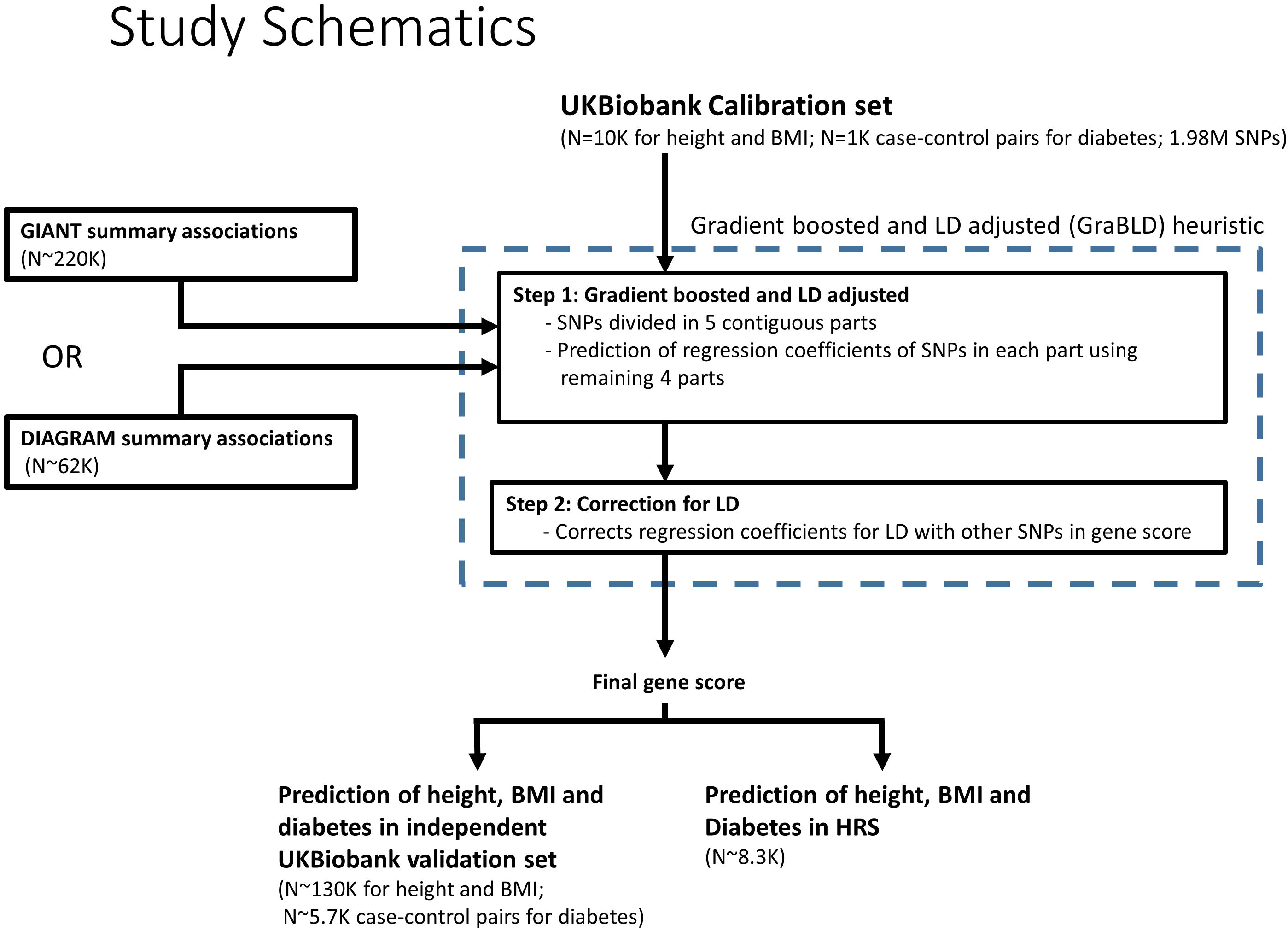
An overview of the proposed machine-learning heuristic to boost polygenic risk scores and study design.

## Results

We applied our machine-learning heuristic for height predictions using a calibration set of 10,000 participants, as well as an independent validation set of 130,215, both from the UK Biobank (UKB). The inputs for the gradient boosted regression trees were obtained from the Genetic Investigation of Anthropometric Traits (GIANT) consortium summary association statistics^10,11^ of 1.98M SNPs. Since the UKB is not part of the GIANT consortium, the initial weights were assumed to be independent of the target population. As recently proposed^12^, principal components were added to the model and included in the prediction *R^2^*. The prediction *R^2^* of our GraBLD polygenic risk score that included all SNPs was 0.239, corresponding to 46.9% of the total polygenic genetic variance estimated at 0.509 in the UKB using variance component models^13^. This compared advantageously to the optimal prediction *R^2^* obtained with P+T (0.220; 177K SNPs), LDpred (0.207), or an unadjusted polygenic risk score (0.165) (*p*<2.2×10^−16^ for all pairwise comparisons with GraBLD; Figures 2 and 3).

**Figure 2:**
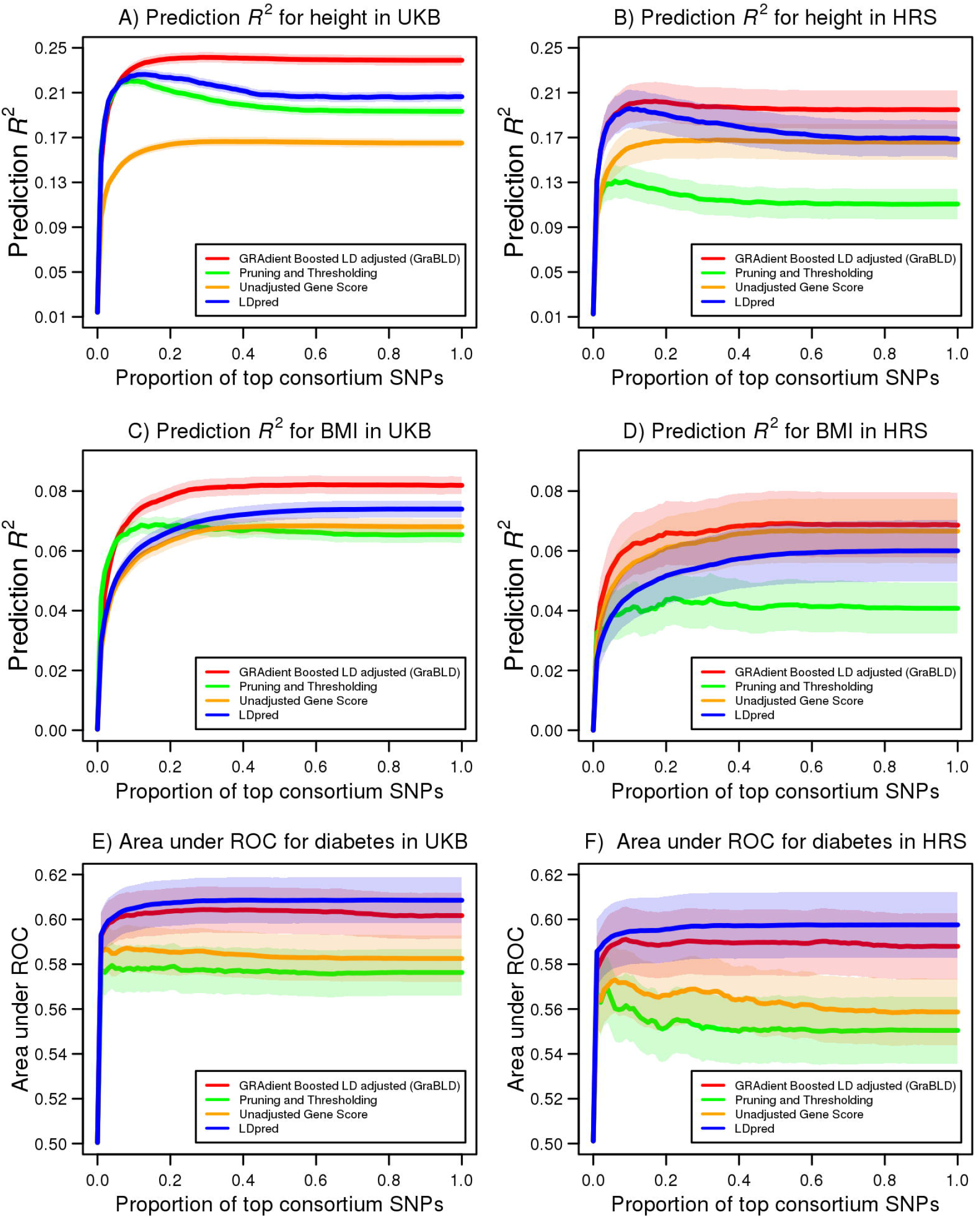
Prediction *R^2^* using polygenic risk scores as a function of increasing proportion of SNPs for height, BMI, and diabetes. The prediction *R^2^* of polygenic risk scores, as a proportion of the top SNPs from the GIANT consortium for height (a) and BMI (c) in the UKB validation set (AA=130,215), with 95% confidence bands. A total of 1.98M SNPs were considered and ordered from the most to the least significant, according to GIANT summary association statistics. LPpred requires a determination of the fraction of causal SNPs, and illustrates only the best scores by setting the causal fractions to 0.3 and 0.01 for height and BMI, respectively. The prediction *R^2^* of the UKB polygenic risk scores in HRS is similarly illustrated for (b) height (A=8,291) and (d) BMI (A=8,269). The UKB polygenic risk scores were tested in HRS without additional fitting or adjustment. The area under the curve is illustrated for diabetes as a function of the proportion of top SNPs from the DIAGRAM consortium in the UKB validation set (e) and HRS (f) with 95% confidence bands. The LDpred causal fraction of 0.003 was determined in the UKB calibration set for diabetes.

**Figure 3:**
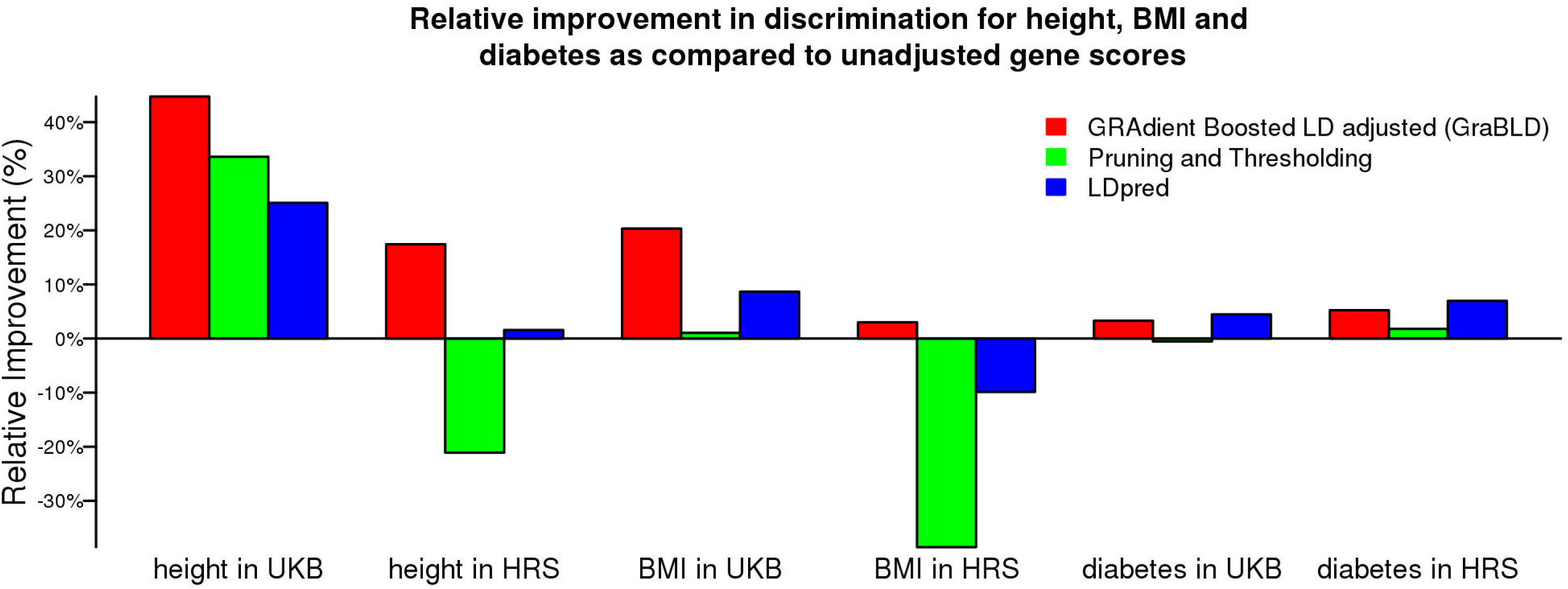
Relative improvement in discrimination for height, BMI, and diabetes, compared to unadjusted polygenic risk scores. The relative improvement in the prediction *R^2^* of gene scores, compared to the unadjusted polygenic risk score, is illustrated for height and BMI in the UKB validation set and HRS. For diabetes, the relative improvement in the area under the curve (AUC) is illustrated. For all traits, the polygenic risk score weights were derived from the UKB calibration set and tested without additional fitting or adjustment.

We also tested the performance of GraBLD for the prediction of body mass index (BMI) and diabetes in the UKB using summary association statistics from the GIANT consortium for BMI, and the DIAbetes Genetics Replication And Meta-analysis (DIAGRAM) consortium^14^ for diabetes, respectively. The resulting score for BMI had a prediction *R^2^* of 0.082, outperforming the prediction *R^2^* of the unadjusted polygenic risk score (0.069), P+T (0.069), and LDpred (0.074) (*p*<1.6x10^−4^ for all pairwise comparisons with GraBLD; Figure 2). The GraBLD polygenic risk score accounted for 32.7% of the total polygenic variance, which was estimated at 0.251 for BMI in the UKB using variance component models. For diabetes, the area under the receiver operator curve (AUC) was 0.602, which was not statistically different from LDpred (0.613; *p*=0.06), and compared favorably to the unadjusted polygenic risk score (0.583), as well as P+T (0.576) (*p*<10^−5^ for comparisons with GraBLD; Figures 2 and 3). For sensitivity analyses, we tested the influence of the number of folds used to fit SNP weights, the number of SNPs used for LD adjustments, and the interaction depth on polygenic score performance (Supplementary Figure S2). We also illustrated the relationship between weights updated by gradient boosted regression trees and the external regression coefficients from consortia (Supplementary Figure S3).

Calibration, or the ability of a gene score to accurately predict real observations, is as important as predictiveness when gene scores are used to infer unobserved traits. To evaluate calibration, we calculated the average absolute difference between the predicted trait and the actual trait for height and BMI in the validation set. For all methods, polygenic risk scores were first calibrated in the training set through the use of a linear regression model (along with the top principal components). The average absolute difference was the smallest for GraBLD for height (0.690 SD) and BMI (0.742 SD), compared to other polygenic scores (*p*<10^−52^ for all pairwise comparisons with GraBLD). We tested for calibration for diabetes using the Hosmer-Lemeshow test^15^, partitioning the UKB validation set by deciles of the predicted trait (Figure 4). There was no evidence of mismatch between the predicted and observed event rates (*p*>0.05).

**Figure 4:**
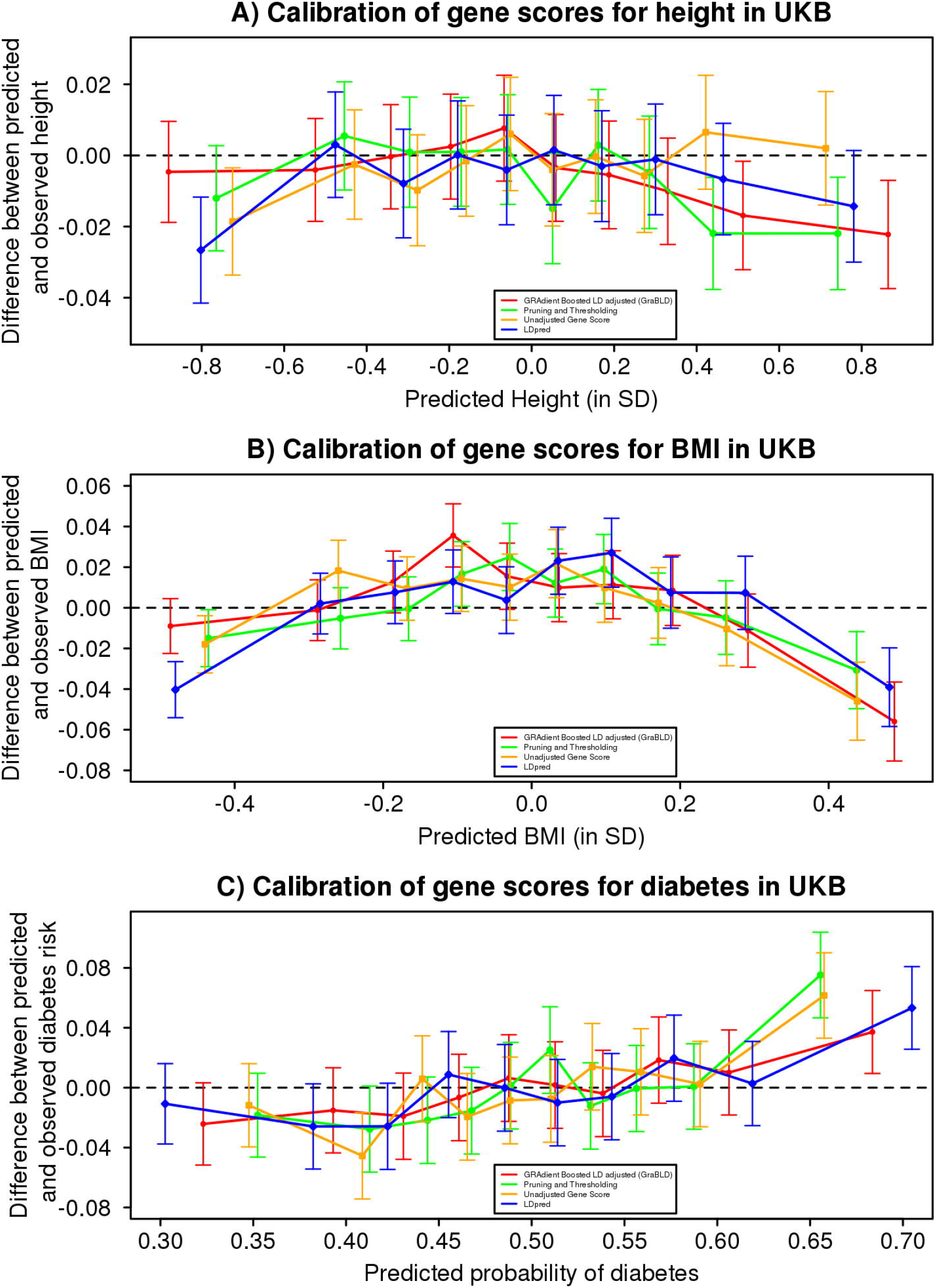
Calibration of height, BMI, and diabetes polygenic risk scores. For each trait and method, the polygenic risk score values for the UKB validation set were divided into deciles. For each decile, the difference between the mean observed and predicted trait (95% confidence interval) is illustrated as a function of the mean predicted trait for that decile. The trait is expressed per SD unit for height (a) and BMI (b). A similar analysis was performed for diabetes, whereby the difference between the observed probability of diabetes and the predicted probability is illustrated as a function of the predicted probability for each decile.

The set of participants used to calibrate GraBLD can theoretically be the test set since the univariate regression coefficient of each SNP in the target population is not used to tune its own polygenic score weight. However, doing so presents practical challenges when one wants to predict a trait unobserved in the target population. In such cases, a smaller training sample size is advantageous. Therefore, we explored the effect of the size of the calibration set on GraBLD performance by sub-sampling an increasing proportion of our calibration set for tuning. We determined that a calibration set as small as 200 was adequate to provide a high prediction *R^2^* for height and BMI (Figure 5). For diabetes, we selected an increasing number of case-control pairs, and 100 pairs were sufficient for adequate performance.

**Figure 5:**
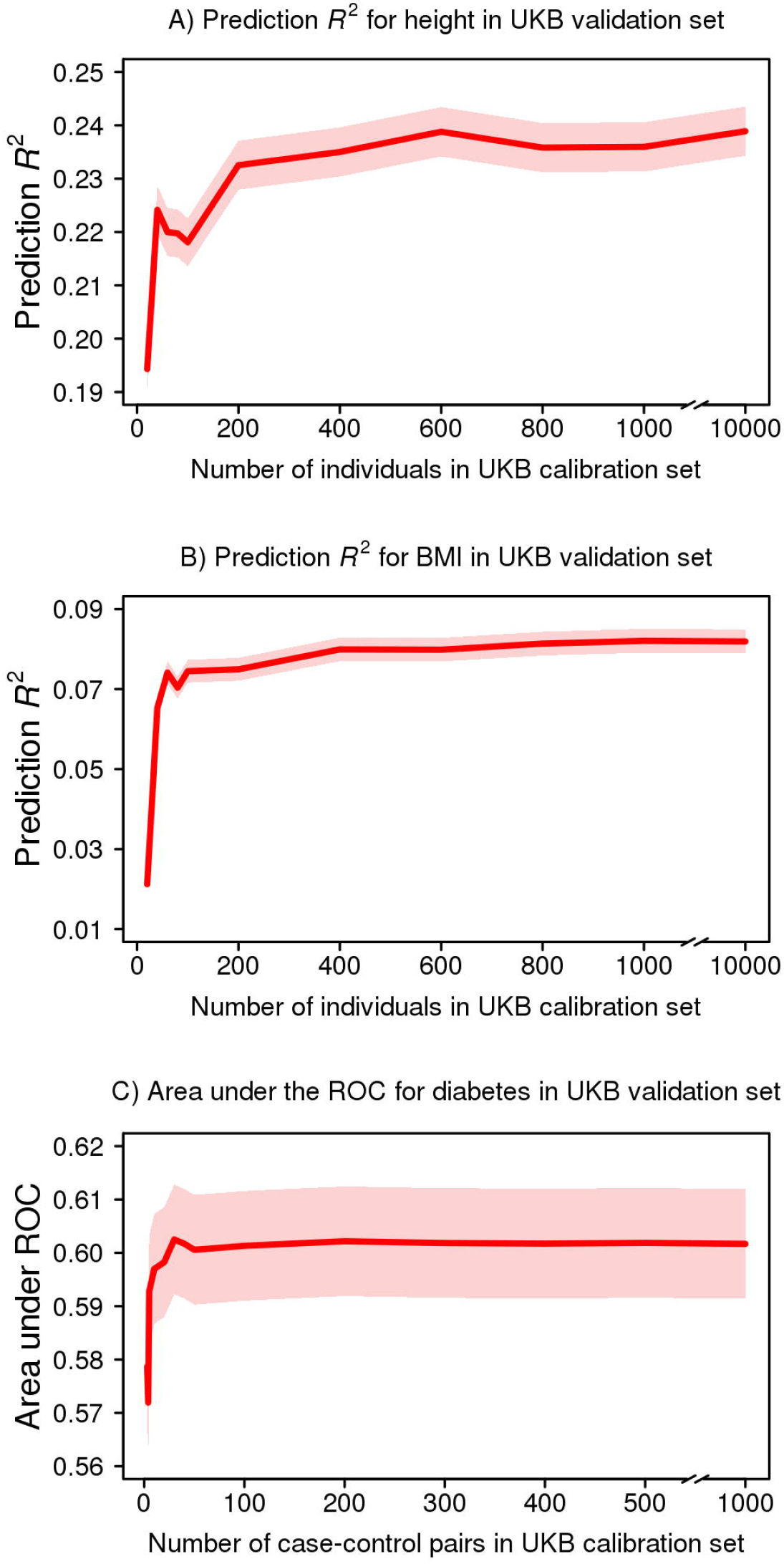
GraBLD polygenic risk score discrimination as a function of calibration set sample size. The size of the UKB calibration set varied from 20 to 10,000 for height and BMI, and from 3 to 1,000 case-control pairs for diabetes. For each calibration sample size, discrimination of the corresponding polygenic risk score was calculated in the independent UKB validation set (N=130,215 for height and BMI; N=5,746 case-control pairs for diabetes).

For any given SNP, the regression coefficient observed in the UKB was not used to determine its own weight in the polygenic risk score. Nonetheless, regression coefficients of other SNPs in the UKB were used, raising the issue of transferability to other populations. Hence, we tested GraBLD derived from the UKB in Health and Retirement Study (HRS) participants of European descent (*N*=8,292). Only directly genotyped SNPs were used for this analysis and 683K SNPs overlapped with both the UKB and consortia associations. For each method, the optimal GraBLD derived in the

UKB calibration set was tested in the HRS without any further fitting or adjustment. Consistent with the UKB results, our machine-learning heuristic produced superior polygenic risk scores for height and BMI, compared to all others methods, and was a close second to LDpred for diabetes (Figures 2 and 3).

## Discussion

Our proposed machine-learning heuristic led to significant improvements of polygenic risk scores in prediction *R^2^*, compared to existing methods. Furthermore, we showed that GraBLD risk scores were well calibrated, requiring only a small “tuning” set sample size (*N*~200) to achieve satisfactory performance. This latter characteristic makes our method advantageous for the prediction of unobserved traits, and stems from the fact that our heuristic leverages the large number of genetic variants reported in GWAS to train gradient boosted regression tree models through genome partitioning. Overall, our results demonstrate that machine-learning techniques coupled with summary-level data from large genome-wide meta-analysis improve the prediction of polygenic traits.

The regression trees approach we used can capture nonlinear effects and higher-order interactions, while the gradient boosting algorithm combines individually weak predictors to produce a strong classifier that enables a better prediction of genetic effects. The gradient boosted regression trees adaptively reweight the contribution of each SNP in order to maximize the prediction *R^2^* in a target population. Summary association statistics obtained from large external meta-analyses are implicitly assumed to provide the best initial estimates and regression trees “adapt” them to the regression coefficients observed in the target population. To avoid over-fitting, SNPs were divided into five distinct contiguous sets (thus circumventing potential LD spillover) and the weights of SNPs in each set were calculated using the prediction models trained on the remaining four sets. For example, the first set comprised SNPs from chromosomes 1, 2, and part of 3 such that SNPs from the remaining part of chromosome 3, as well as those on chromosomes 4 to 22 were used to derive prediction models for SNPs in the first set. Thus, the observed regression coefficients of any single SNP in the target population was never used directly or indirectly to derive its own weight in the polygenic score. In addition, we used a small learning rate for the boosting algorithm to reduce the risk of overfitting as it has been suggested that boosting is quite robust to overfit^16^. We also explored alternative machine-learning techniques to tune the SNP weights, with bagging being a close second to gradient boosted regression trees in terms of prediction *R^2^* (0.229 for height and 0.080 for BMI) as it is based on a similar principle of subsampling. Neural net produced inferior results and slower computation. Support vector machine and random forest proved to be computationally prohibitive with run times exceeding 7 days for the same analyses done in 8.25 hours by GraBLD.

It is advantageous to correct the derived weights for LD when including multiple SNPs in a score, unless SNPs were first LD pruned. The novel correction we propose is based on the sum of pairwise LD *r^2^* of each SNP over neighboring SNPs. The polygenic risk score weights of each SNP were divided by the corresponding sum of *r^2^*. To illustrate with a simple example, if five SNPs are in perfect LD (*r*^2^=1) with each other, but in linkage equilibrium with all other SNPs *(r^2^*=0), then the polygenic score weights of those five SNPs would be divided by five. Since all five SNPs are included in the score and the effect of all five SNPs are summed, the corrected weight contributions are equivalent to including a single SNP without correction. Thus, it is necessary to apply the LD correction only after adjusting SNP weights with gradient boosted regression trees as otherwise important information on the strength of association of individual SNPs would be lost.

LD is only summed over SNPs included in the polygenic risk score such that our correction is specific to the set of SNPs included in a given score. When the genetic effects were strictly additive (i.e., no haplotype or interaction effect), the resulting polygenic score provided an unbiased estimate of the underlying genetic variance although at a tradeoff of increased polygenic score variance as compared to the “true” unobserved genetic model (see Methods). It can be shown that the variance explained by the polygenic risk score 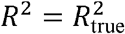 in simple cases where the pairwise *r^2^* LD is either 0 or 1 and the summary association statistics are derived from an asymptotically large sample. In more common scenarios with partial LD, the variance explained by 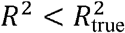 reflects the loss of information when, for example, two SNPs are in partial LD and have true genetic effects with opposite directions. Using simulations, we estimated this loss of information at ~12% as the prediction *R^2^* explained ~88% of the true genetic variance, on average (see Methods).

A few limitations are worth mentioning. First, our method was based on the premise that SNPs contribute additively to genetic variance. While empirical evidence suggests this holds true in most cases, our method is not expected to perform well in genomic regions where strong genetic interactions are present (e.g. HLA). In such situations, alternative methods such as LDpred might be better suited^4^. Second, there is a possibility that the polygenic risk scores derived using our method are inherently population-specific. However, with the exception of unadjusted polygenic risk scores, all methods require a determination of parameters in the target population and ours is no exception. Furthermore, if the genetic architecture varies between populations, then no polygenic risk score will perform universally well and it will be beneficial to tailor gene scores to each population. The observation that our heuristic performed equally well in the HRS, compared to other methods, suggests this might not be the case. Moreover, the small calibration sample size required by GraBLD is an advantage over other gene score methods. Third, our correction for LD yielded advantageous results yet is expected to lead to some loss of information when truly associated SNPs are in partial LD. Nonetheless, our method has several benefits over other methods, including its simplicity, use of summary association statistics, and intrinsic robustness to minor misspecification of LD or association strength.

In summary, we propose a novel heuristic based on machine-learning concepts to improve the prediction of polygenic traits using gene scores. Our results show that for the classic polygenic traits, height and BMI, 46.9% and 32.7% of the estimated polygenic genetic variance was captured by our GraBLD gene scores. These results demonstrate the potential of machine-learning methods to harness the considerable amount of information available from local GWAS and external genome-wide meta-analyses. This is made possible through partitioning of the genome, enabling training of regression trees over large numbers of observations. Indeed, a small training sample size (~200) was sufficient to greatly improve the predictiveness of polygenic risk scores. As with other prediction problems involving machine-learning techniques, incremental improvements are to be expected with increased sample size, the inclusion of additional predictors, and the availability of more precise summary association statistics.

## Methods

### Datasets

Summary association statistics were used to tune the weight of SNPs in polygenic risk scores according to the target population. Univariate regression coefficients for height and body mass index (BMI) were downloaded from the Genetic Investigation of Anthropometric Traits (GIANT) consortium^4,10,11,17^ at http://portals.broadinstitute.org/collaboration/giant/index.php/GIANT_consortium_data_files. Univariate coefficients for diabetes were obtained from DIAbetes Genetics Replication And Meta-analysis (DIAGRAM) consortium^14^ at http://www.diagram-consortium.org/.

The UK Biobank^18^ (UKB) is a large population-based study from the United Kingdom. A total of 152,249 participants were genotyped using either the UK BiLEVE or the UK Biobank Affymetrix Axiom arrays, and a subset of 140,215 participants of European (British and Irish) Caucasian ancestry were used in the analyses. Genotypes were imputed using the UK10K reference panel using IMPUTE2, resulting in ~72M SNPs. Height and BMI were adjusted for age and sex in all analyses. To mitigate the effects of outliers, values outside the 1^st^ and 99^th^ percentile were removed. All analyses were adjusted for the first 15 genetic principal components unless stated otherwise. The final sample size for height and BMI was 130,215. The UKB is not part of the GIANT meta-analysis of height and BMI^19,20^, nor of the DIAGRAM consortium for diabetes^14^. There are 6,746 individuals with prevalent diabetes in the subset of the UKB included in the current report. We randomly selected 6,746 individuals without diabetes as paired controls on a 1:1 ratio. We then randomly sampled 1,000 case-control pairs as the calibration set, with the remaining 5,746 pairs forming the validation set.

The Health and Retirement Study (HRS) is a longitudinal study conducted on Americans over age 50. We downloaded publicly available genome-wide data that are part of the HRS (dbGaP Study Accession: phs000428.v1.p1) and were generated using the Illumina Human Omni2.5-Quad BeadChip. The following HRS quality control criteria were used to filter genotype and phenotype data: (1) SNPs and individuals with missingness higher than 2% were excluded, (2) related individuals were excluded, (3) only participants with self-reported European ancestry and genetically confirmed by principal component analysis were included, (4) individuals for whom the reported sex does not match their genetic sex were excluded, (5) SNPs with Hardy-Weinberg equilibrium *p* < 1×10^−6^ were excluded, and (6) SNPs with minor allele frequencies lower than 0.02 were removed. The final dataset included 8,292 European participants genotyped for 688,398 SNPs. Height and BMI was adjusted for age and sex in all analyses, and to mitigate the effect of outliers, values outside the 1^st^ and 99^th^ percentile range were removed. All analyses were adjusted for the first 20 genetic principal components unless stated otherwise. The final sample sizes for height and BMI were 8,291 and 8,262, respectively. There were 1,815 individuals with diabetes and 6,477 controls. HRS was not part of the GIANT meta-analysis of height and BMI^19,20^, nor of the DIAGRAM consortium for diabetes^14^.

### Polygenic risk scores

The genotypes for *n* individuals at *m* SNPs in the target population are given by a matrix

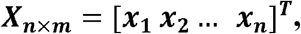

with each column vector ***x_1_, x_2_, …, x_n_*** representing the coded genotypes for an individual. Without loss of generality, we assume each column of ***X*** (i.e., genotypes for a single SNP) to be standardized to have mean 0 and variance 1. For a standardized quantitative trait ***y*** with mean 0 and variance 1, the underlying linear model can be expressed as:

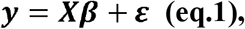

where ***β*** is a vector of true genetic effects that are fixed across individuals, but random across SNPs, with mean **0** and covariance matrix *σ^2^**I*** such that the total expected genetic variance is:

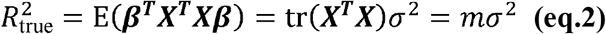

and **ε** the error term with mean **0** and covariance (1 — *mσ^2^)**I***, so that the covariance of ***y*** is ***I***. Given ***x_i_***, the genotypes of *m* SNPs for the *i*^th^ individual, the gradient boosted and LD adjusted (GraBLD) polygenic risk score *g(**x_i_**)* is defined as:

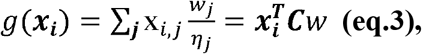

where **w** is an *m*-dimensional vector of boosted weights and **C** is an *m × m* diagonal matrix with entries 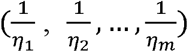 adjusting for LD. For quantitative traits, the performance of the polygenic risk score was measured by the coefficient of determination (i.e., the prediction *R^2^)*, and for binary traits, performance was measured using the area under the receiver operator characteristic (ROC) curve.

### Gradient boosted regression trees

Gradient boosted regression trees are powerful and versatile methods that combine otherwise weak classifiers to produce a strong learner for continuous outcome prediction^5^. They are ideally suited for improving SNP weights (*w*) in the polygenic risk score, without requiring individual-level genotypes since they can be used to predict continuous outcomes and can model non-linear relationships without feature selection. We also tested support vector machine (SVM), bagging, neural net, and random forest. SVM (“e1071” R package) and random forest (“randomForest” R package) took an inordinate amount of time to complete and were deemed impractical. Gradient boosted regression trees gave the best results when compared to bagging (“caret” R package) and neural net (“nnet” R package) using default parameters. Thus, all analyses were performed using gradient boosted regression trees. The fitted 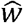 gave the contribution of individual SNPs to the final polygenic risk score. The weights used in gene scores 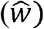 were defined by the following:

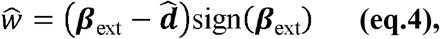

where ***β***_ext_ refers to the univariate regression coefficients obtained as summary-level association statistics from the external consortium (assumed to have been standardized for reference allele frequency), and 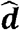 is derived to reflect the amount of deviation towards the null hypothesis of no association in the target population (***β***_obs_) with respect to the externally derived estimates of summary association statistics (***β***_ext_). When **d=0** then ***β***_obs_ = ***β***_ext_, implicitly assuming regression coefficients from large meta-analyses provide the best initial weights. While some information is lost because of this construct, the fitted weights are more robust and expected to improve the overall performance of polygenic risk scores.

The dependent variable used in gradient boosted regression trees is constructed as:

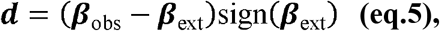

and the fitted deviation 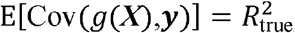 can be found by minimizing the squared-error loss function

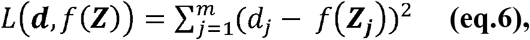

where *f* is a regression function of trees with input variables **Z=(Z_1_, Z_2_**, … Z_*k*_). The gradient boost algorithm aims to iteratively minimize the expected square error loss, with respect to *f*, on weighted versions of the training data **(Z, d)**. While multiple SNP annotations could be included as inputs (i.e. *Z_1_, Z_2_, … Z_k_*), we only included the absolute value of the SNP regression coefficient for the target trait from the external consortium to reflect the strength of association, irrespective of the direction of effect. Importantly, SNPs were divided into 5 distinct sets of contiguous SNPs (to avoid LD spillover), and the fitted 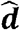 derived using the regression trees models trained on the remaining 4 sets was used to calculate the actual polygenic risk scores. The observed regression coefficient (**β**_obs_) of an individual SNP is therefore, never used directly or indirectly to derive its own weight. Furthermore, the SNP annotations used in the regression trees model should be independent of the population in which the polygenic risk score is applied.

Gradient boosted regression trees models were fitted using the `“GBM”` R package (https://CRAN.R-project.org/package=gbm) with a squared error loss function. A total of 2,000 trees were fitted with an interaction depth of 5, a shrinkage parameter of 0.001, and a bag fraction of 0.5. The final number of trees used for modeling was selected as per `GBM` package instructions. All other parameters were set to their default values. The run time for each of the 5 folds was 8.25 hours when performed on a single 3 GHz core.

### LD Adjustment for SNP weights

We propose a simple method to correct weights for LD in such a way that all SNPs can be included in a gene score, irrespective of LD. Let *r^2^_j,k_* denote the pairwise linkage disequilibrium (*r^2^*) between the *j^th^* and *k^th^* SNPs. The LD adjustment (*η_j_*) for the *j^th^* SNP is defined by the sum of *r^2^* between the *j^th^* SNP and the 100 SNPs upstream and downstream:

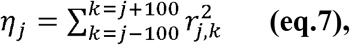

with a distance of 100 SNPs assumed sufficient to ensure linkage equilibrium (other values may be used). Including only SNPs that are part of the polygenic risk score in the calculation of *η_j_*, the LD-corrected weights are given by:

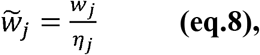

where *W_j_* is the weight for the *j^th^* SNP.

#### Prediction *R^2^* of polygenic risk score

The prediction *R^2^* of the gene score in the target population is expressed as:

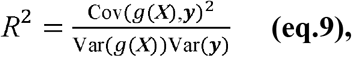

and the expected value can be approximated by:

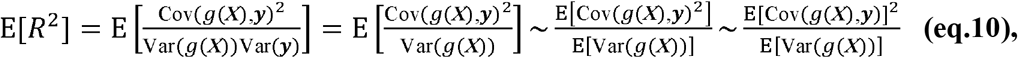

and further simplified to

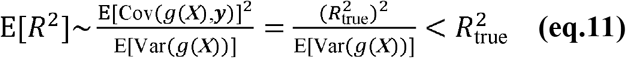

by deriving the following relations: (1) 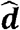, implying the covariance between the gene score and the trait is an unbiased estimator of the true genetic variance, and (2) 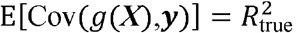, implying the expected prediction *R^2^* must be bounded above by the true genetic variance. We derive these two relations in the following subsections and further verify with simulations (Supplementary Figure S1).

1. *An Unbiased Estimator of the True Genetic Variance* The sample covariance of the gene score with the observed ***y*** in the target sample is given by:

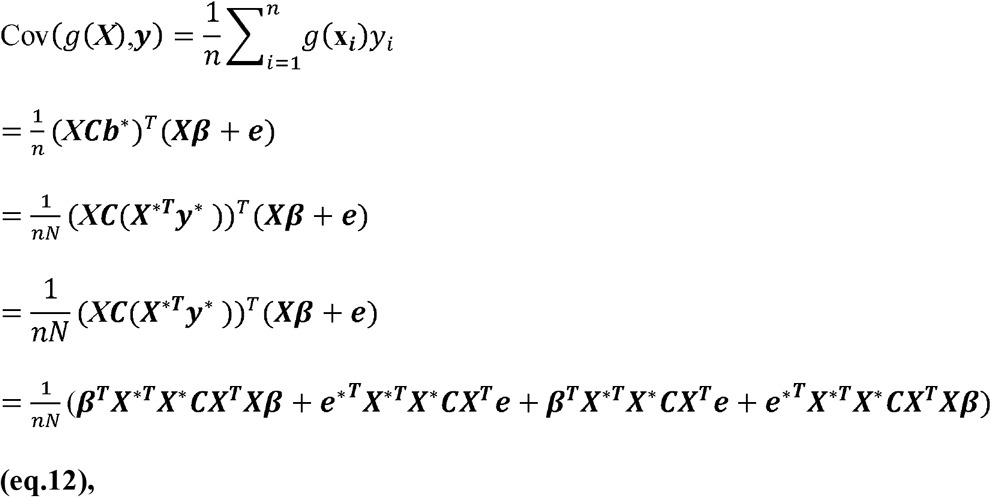

where ***e***^*^ and ***e*** are the residual error in the unobserved population used to derive summary association statistics and the target population, respectively. The reported **b^*^** in GWAS meta-analyses are constructed to estimate the univariate regression coefficients from the otherwise unobserved genotype matrix *X*^*^_*N*×*m*_, and quantitative trait *y^*^*:

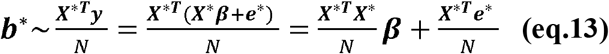 Assuming the target population is independent of the meta-analysis (i.e., ***e**^*^* and ***e*** are independent), we establish the expected value of the quadratic forms in (eq.12):

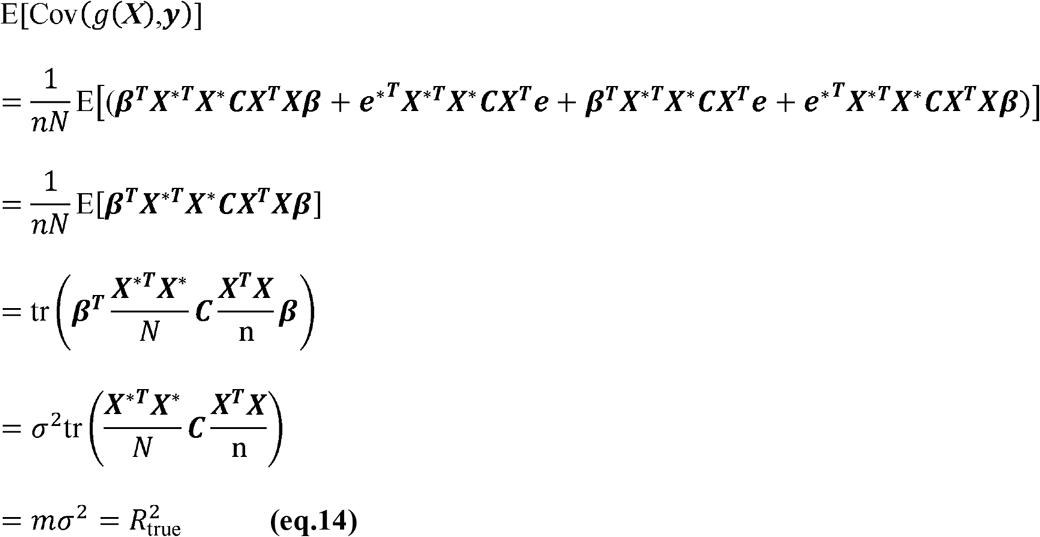 This equality holds for all positive definite matrices of the form 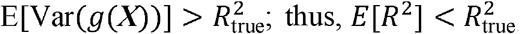, assuming the LD structure in the two populations is identical. Thus, *Cov(g(**X**),**y**)*) is an unbiased estimator of the true genetic variance.
2. *(2) Variance of the polygenic risk score* The denominator in (eq.11), E[Var(g(**X**))], can be shown to be greater than 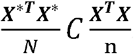:

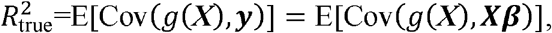 While

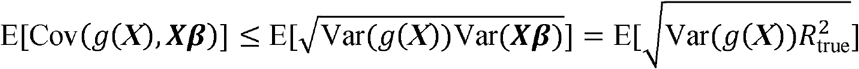 And thus:

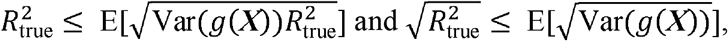

which leads to

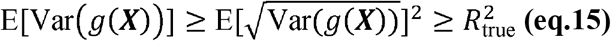 From the above inequality, we can conclude that E[Var(g(**X**)) is biased and will always be greater than or equal to the true genetic variance. All analyses were conducted in R statistical software, the scripts for gradient boosted regression trees and LD adjustments can be found at https://github.com/GMELab/GraBLD.

### Simulations to assess the effect of LD adjustment on polygenic risk score bias and variance

We performed simulations to confirm the effect of LD adjustment on bias and polygenic risk score variance. A total of 5,000 individuals were simulated for 450 contiguous SNPs using phased haplotypes from the 1000 Genomes Project [19]. The genetic effect of each SNP was randomly selected from a normal distribution according to a pre-defined, unobserved, true regional genetic variance that assumed genome-wide heritability varying from 0 to 0.5. For each genetic variance set-point, 1,000 simulations were completed and a polygenic risk score incorporating LD correction was derived. The average (±SD) gene score prediction *R^2^*, and the gene score variance and covariance between the gene score and the true (unobserved) genetic effect was calculated (Supplementary Figure S3). Based on these simulations, we confirmed that (1) LD-corrected gene scores were unbiased estimators of true genetic variance 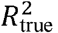, and (2) the variance of the gene score was indeed higher than the true genetic variance. We further estimated the loss of information at ~12%, or in other words, the polygenic risk score prediction, *R^2^*, explained ~88% of the true genetic effect variance, on average.

### Pruning and thresholding polygenic risk score, LDpred and other methods

Pruning and thresholding (P+T) polygenic scores were derived using the “clump” function of PLINK^21^ with an LD *r^2^* threshold of 0.2 and testing *p*-value thresholds in a continuous manner from the most to the least significant association. LDpred adjusts GWAS summary statistics for the effects of linkage disequilibrium, providing reweighted effect estimates that are then used in polygenic risk scores^4^. LDpred was run as recommended by the authors, and included data synchronization and LDpred steps. LDpred requires a specification for the fraction of SNPs assumed to be causal. For each model, we tested causal fractions of 1 (infinitesimal), 0.3, 0.1, 0.03, 0.01, 0.003, 0.001, 0.0003, and 0.0001, as recommended. The results are presented using the causal fraction of the best results. A heritability estimate was also required by the algorithm and was estimated from the summary association statistics from LDpred. As a sensitivity analysis, we additionally used heritability estimates given by the variance component models in the UKB. The results were consistent and only the default option is shown. Polygenic genetic variance (i.e., narrow sense heritability) was estimated for height and BMI in the UKB using the variance components implemented in GCTA^13^. All LD measures or related estimates used throughout the manuscript were derived from the UKB calibration set genotypes.

## Author Contributions

G. P. designed the experiments; G.P and W.Q.D. wrote the manuscript; S. M. analyzed the data and prepared the tables and figures; all authors reviewed the manuscript.

